# Spatial proteomics and transcriptomics of placenta accreta spectrum

**DOI:** 10.1101/2024.03.21.585167

**Authors:** Helena C Bartels, Sodiq Hameed, Constance Young, Myriam Nabhan, Paul Downey, Kathleen M Curran, Janet McCormack, Aurelie Fabre, Walter Kolch, Vadim Zhernovkov, Donal J Brennan

## Abstract

In severe Placenta Accreta Spectrum (PAS), trophoblasts gain deep access in the myometrium (placenta increta). This study investigated alterations at the fetal-maternal interface in PAS cases using a systems biology approach consisting of immunohistochemistry, spatial transcriptomics and proteomics. We identified spatial variation in the distribution of CD4^+^, CD3^+^ and CD8^+^ T-cells at the maternal-interface in placenta increta cases. Spatial transcriptomics identified transcription factors involved in promotion of trophoblast invasion such as AP-1 subunits ATF-3 and JUN, and NFKB were upregulated in regions with deep myometrial invasion. Pathway analysis of differentially expressed genes demonstrated that degradation of extracellular matrix (ECM) and class 1 MHC protein were increased in increta regions, suggesting local tissue injury and immune suppression. Spatial proteomics demonstrated that increta regions were characterised by excessive trophoblastic proliferation in an immunosuppressive environment. Expression of inhibitors of apoptosis such as BCL-2 and fibronectin were increased, while CTLA-4 was decreased and increased expression of PD-L1, PD-L2 and CD14 macrophages. Additionally, CD44, which is a ligand of fibronectin that promotes trophoblast invasion and cell adhesion was also increased in increta regions. We subsequently examined ligand receptor interactions enriched in increta regions, with interactions with ITGβ1, including with fibronectin and ADAMS, emerging as central in increta. These ITGβ1 ligand interactions are involved in activation of epithelial–mesenchymal transition and remodelling of ECM suggesting a more invasive trophoblast phenotype. In PAS, we suggest this is driven by fibronectin via AP-1 signalling, likely as a secondary response to myometrial scarring. Overall, this study suggests the biological processes leading to deep trophoblast invasion in the myometrium in placenta increta are as a result of upregulation of transcription factors and subsequent genes and proteins which promote trophoblast invasion. This occurs in a locally immune suppressed environment, with increased ECM degradation suggesting these findings are secondary to iatrogenic uterine injury.

**Significance statement:** Placenta Accreta Spectrum (PAS) is a rare pregnancy complication, where the placenta fails to separate from the womb resulting in severe bleeding, which is associated with significant maternal morbidity and mortality. As Caesarean section rates increase, the incidence of PAS is increasing. The underlying pathophysiology of PAS is poorly understood. Here, we apply a spatial multi-omic approach to explore the biologic changes at the maternal-fetal interface in severe PAS (placenta increta). Using spatial transcriptomics and proteomics, we identified genes and proteins that are dysregulated in severe PAS involving processes such as extracellular matrix degradation, local immune suppression and promotion of epithelial–mesenchymal transition. This study provides new insights into the biological changes and underlying pathophysiology leading to placenta increta.

## Background

Placenta Accreta Spectrum (PAS) is a severe pregnancy complication associated with significant maternal morbidity and mortality (1, 2). The incidence of PAS is increasing, largely related to the increase in the number of women giving birth by Caesarean section (3, 4). In PAS, the placenta has abnormally adhered to or invaded the myometrium, and fails to separate from the uterus after the birth, resulting in massive haemorrhage (5, 6).

The pathophysiology of PAS is controversial. However, it is generally agreed that fetal trophoblast cells are in direct contact with myometrium due to an absent or deficient decidua as a result of prior uterine injury (5, 7). More severe cases are categorised based on deep infiltration of the myometrium by trophoblast cells (7). Extravillous trophoblast cells (EVT) exhibit sustained proliferative signalling and an overly invasive phenotype in PAS (8, 9). Single cell RNA-Seq studies in PAS identified upregulation of genes promoting invasion of EVTs, with one study describing a distinct excessive invasive cytotrophoblast cell types (LAMB4+ and KRT6A+) (10). Several genes involved in promoting EVT invasion, such as PRG2 and AQPEP, were abnormally expressed in the trophoblasts and fetal membranes of PAS cases (11). A further study found negative regulators of cell migration, such as podocin and apolipoprotein D, were downregulated in PAS, suggesting a pro-adhesive signature (12). Immune changes at the maternal-fetal interface in PAS, particularly in the absence of a decidua, may also facilitate excessive trophoblast invasion as demonstrated by reduced populations of CD4^+^ T cells and uterine natural killer cells (13–15).

These studies suggest that there are biological changes in the placenta in PAS cases, where trophoblasts express an overly invasion phenotype which may be facilitated by a focal immunosuppressive microenvironment. These changes may be secondary to a defective decidua as a result of primary uterine injury when the normal regulators of trophoblast invasion are absent (16).

Furthermore, PAS cases are often associated with increased angiogenesis at the uteroplacental interface (17). This results from EVTs gaining access to and remodelling deep maternal spiral arteries as a result of absent or defective decidualisation and scar dehiscence (18). Consequently, there is chronic high velocity blood flow into the intervillous space, resulting in an excess deposition of fibrinoid at the maternal-fetal interface in PAS (19). Placental fibrinoids are important in normal placentation for creating a glue effect to attach the placenta to the myometrium (20), hence the excessive deposition in PAS is thought to contribute to the abnormal placental attachment in PAS (19).

We hypothesise that the biologic changes at the maternal-fetal interface seen in severe PAS represent an abnormal upregulation of the normal processes required for placental invasion as a result of defective decidualisation from iatrogenic uterine injury. We investigate this hypothesis by performing immunohistochemistry, spatial transcriptomics and proteomics of the maternal-fetal interface in PAS, and comparing our results to publicly available single cell RNA-seq data. The experimental workflow utilised in this study is summarised in Figure 1.

**Figure 1:**
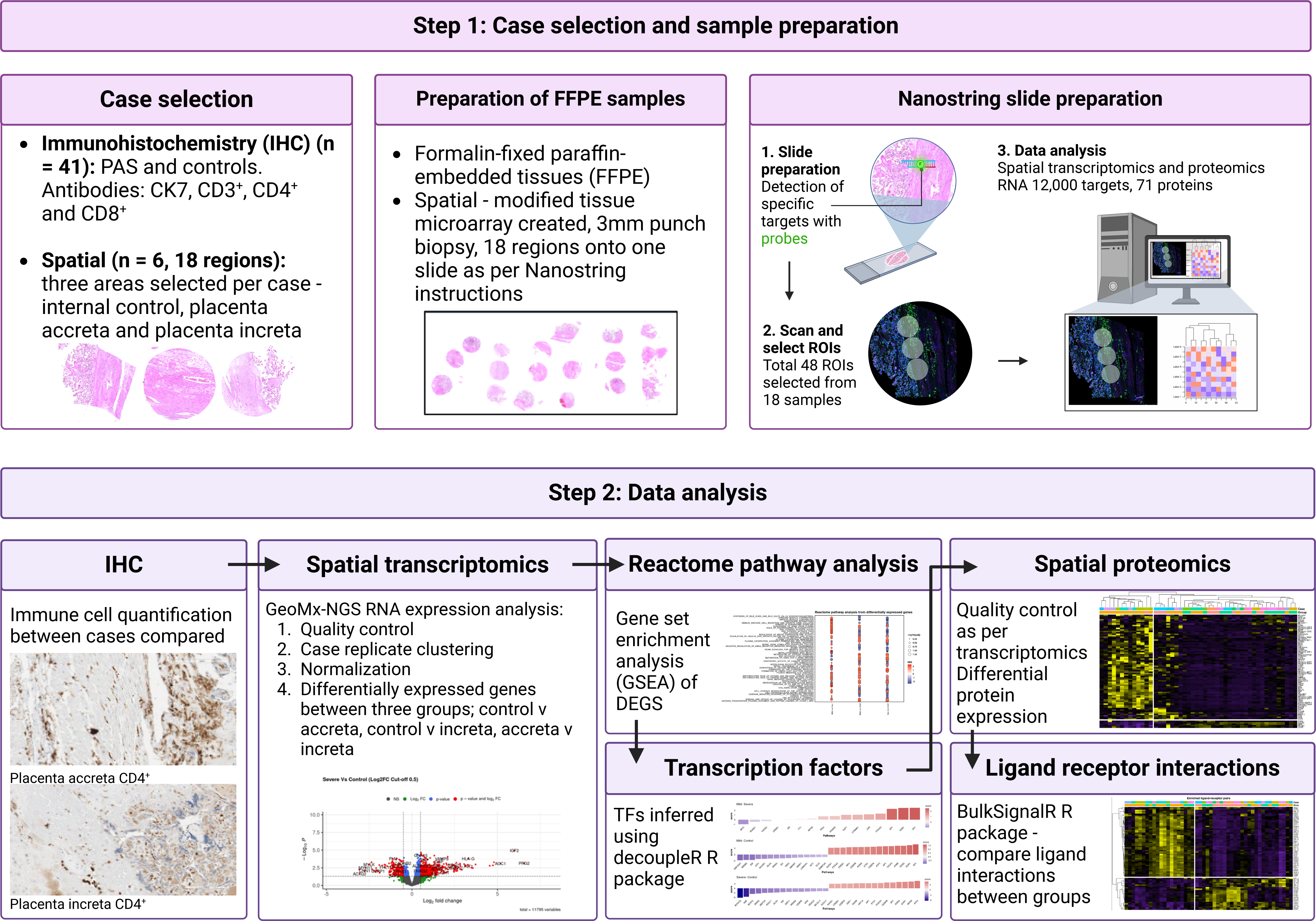
Experimental workflow summary.

## Results

### Differences in immune cell infiltrate at the fetal-maternal interface

We used immunohistochemistry to examine the relationship between the degree of placental invasion (as defined by FIGO grading and Society of Paediatric pathology (5, 7), methods) and immune cell infiltrate at the maternal-fetal interface (Supplementary Figure 1, 2). There was a dense immune infiltrate of CD3^+^, CD4^+^ and CD8^+^ T cells in the placenta accreta cases (n=18) at the invasive front of fetal trophoblast cells (represented by CK7 expression) which were in direct contact with maternal myometrium (Figure 2a), however in placenta increta cases (n=17), we noted significant heterogeneity in the location and number of immune cells at the maternal-fetal interface (Figure 2a). In some placenta increta cases, very few immune cells were identified, while others demonstrated a moderate immune infiltrate, although dense clustering as seen in the placenta accreta cases was absent. While the overall T cell density did not differ between control cases (placenta previa with a previous Caesarean section) as demonstrated by CD3^+^ expression, CD4^+^ T cell infiltration was significantly lower in the placenta increta compared to the placenta accreta (p = 0.012) and controls (p = 0.007), (Figure 2b), consistent with a previous study (14). No differences were seen in the number of CD8^+^ T cells. These findings suggested there were differences in the spatial distribution of immune cells in PAS cases, and in order to further explore these differences, we performed spatial proteomics and transcriptomics on a subset of cases.

**Figure 2:**
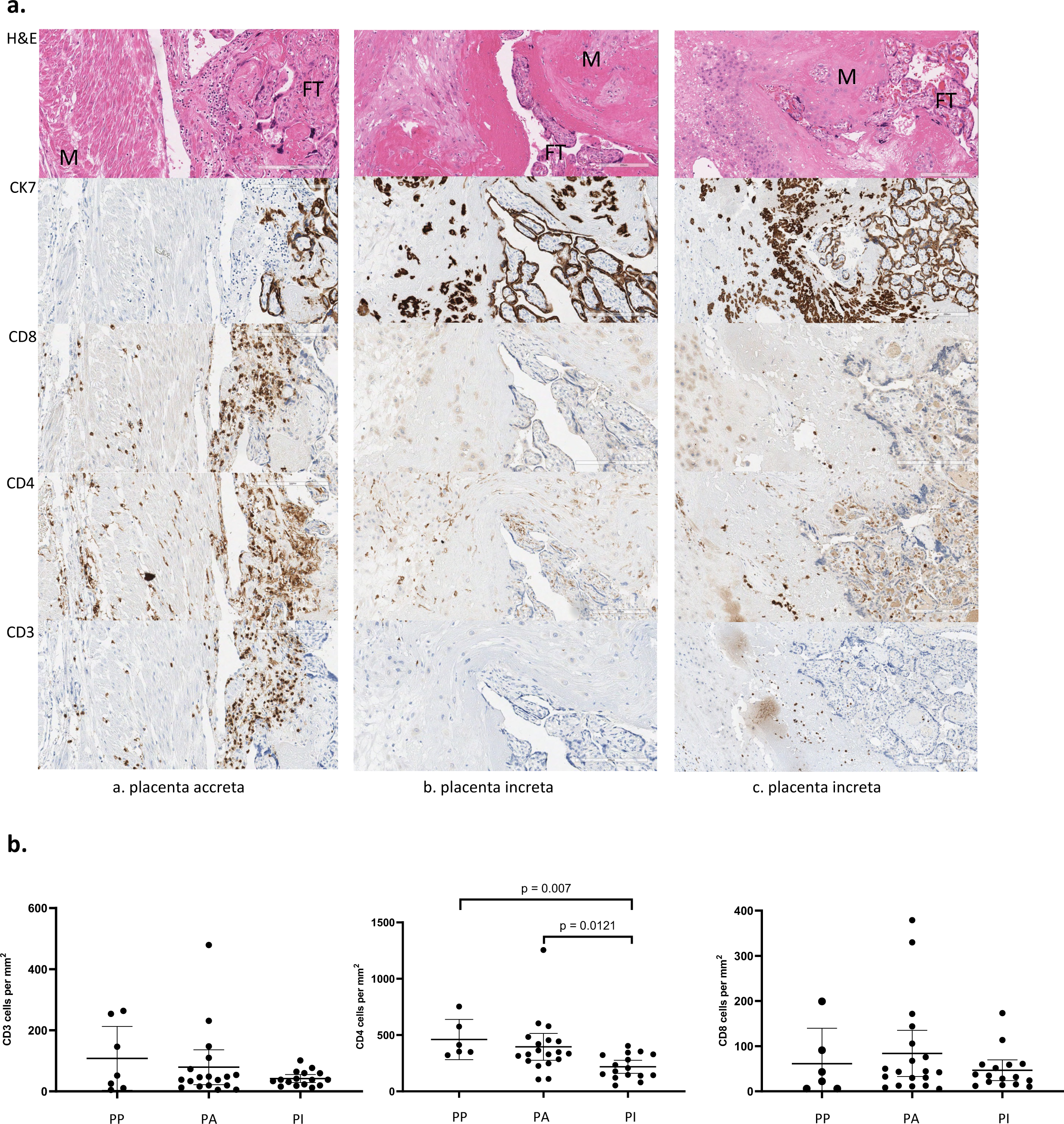
Immunohistochemistry. **a:** Comparison of immune infiltrate between control placenta accreta, and two placenta increta cases. Top row H&E stained slides at 200um magnification showing maternal-fetal interface. Staining for four antibodies; CK7, CD8^+^, CD4^+^ and CD3^+^. CK7 stains for fetal trophoblast (FT) cells; in placenta accreta, FT are in direct contact with the myometrium (M) with no intervening decidua. In placenta increta, FT are seen deeper within the myometrium, and there are thick depositions of fibrinoid (pink). CD8, CD4, and CD3 staining shows clusters of immune cells in the region where FT are in direct contact with the maternal myometrium, no invasion of FT seen beyond the cluster of immune cells. Placenta increta (b) shows minimal immune cells in an area where deep trophoblast invasion is present. In case (c), some immune cells are seen, however these are more widely spread compared to case placenta accreta. **b**: immune cell quantification compared between controls, placenta accreta and placenta increta PAS cases. CD4^+^ cells were significantly reduced in placenta accreta and placenta increta compared to controls. *FT: Fetal Trophoblast. M: myometrium. PA: Placenta Accreta. PI: Placenta Increta. All immunohistochemistry slides at 200um magnification*.

### Spatial transcriptomics: differential gene expression in regions of placenta increta

For spatial transcriptomics, we used the Nanostring GeoMx Digital Spatial Profiling (DSP) (Seattle, WA, USA) platform to detect approximately 12000 targets at the maternal-fetal interface. The number of differentially expressed genes comparing regions of placenta accreta vs. increta, accreta vs. control and increta vs. control was n=16, n=603 and n=947, respectively (Figure 3a). Although we only identified 16 differentially expressed genes between placenta accreta and increta regions, one of these was NF-κB2 which is a regulator of normal trophoblast invasion (21). NF-κB is expressed in syncytiotrophoblast and the stroma of placental villi, and increased protein expression of NF-κB has been described in PAS (22). ALX1, which has been linked to increased epithelial–mesenchymal transition (EMT) and invasion in malignancy (23), was also increased in placenta increta compared to accreta regions. This suggests that while there are only a small number of differentially expressed genes between increta and accreta cases, increta cases demonstrate a more invasive phenotype.

**Figure 3:**
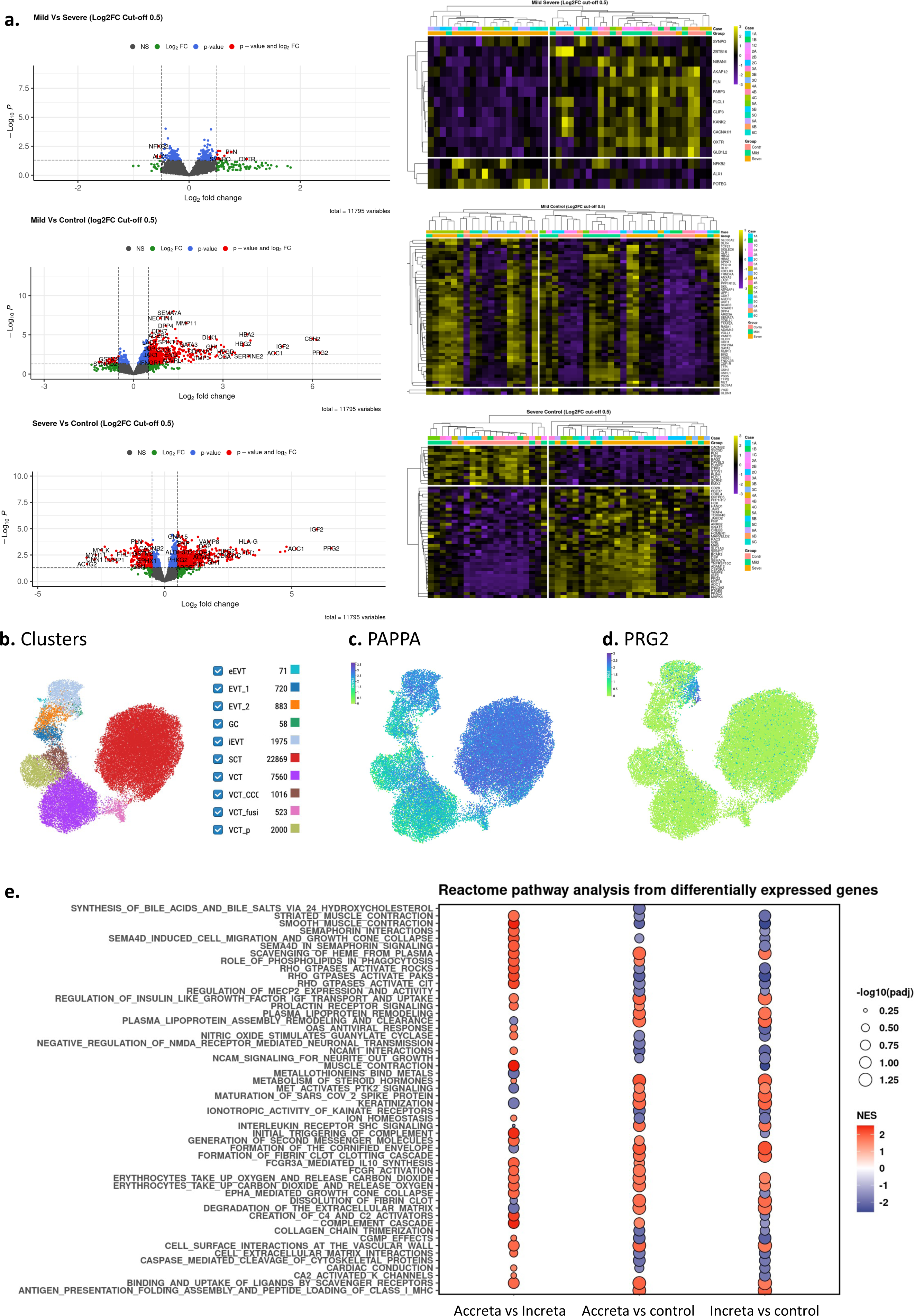
Spatial transcriptomics and gene set enrichment analysis. Figure 3 shows the differentially expressed genes (DEGs) between accreta vs increta, accreta vs control and increta vs control. **3a** Heatmap and volcano plot of differentially expressed genes (DEGS). The number of DEGs for each comparison was n=16, n=603, and n=947 for accreta vs increta, accreta vs control and increta vs control respectively. **3b** Spatially resolved single cell data from maternal-fetal interface available at https://reproductivecellatlas.org) from a hysterectomy specimen - cell clusters from snRNA-seq data, **c** PAPPA expression shown in SCT and in placental giant cells, **d** PRG2 expression iEVTs. **3e** Gene set enrichment analysis of differentially expressed genes; the pathways differentially expressed in each comparison of the DEGs. *eEVT: endovascular trophoblast cells. GC: placental bed giant cells. iEVT: interstitial extravillous trophoblast. SCT: villous syncytiotrophoblast. VCT: villous cytotrophoblast. VCT_CCC: villous cytotrophoblast cytotrophoblast cell columns*.

Comparing placenta increta to control regions, several genes involved in normal trophoblast invasion were identified and had increased expression including Pregnancy Associated Plasma Protein A (PAPPA), PRG2 and IGF-2 (Figure 3a). PRG2 and IGF2 were the two most differentially expressed genes when comparing accreta to control and increta to control regions. PRG2 overexpression has been previously described in PAS (11, 24). PAPPA is a metalloproteinase which acts as a growth promoter in many tissues through release of IGF (25). In normal placentation, EVTs fuse into placental bed giant cells at the maternal interface, where they upregulate the PGR2–PAPPA complex (21), suggesting this interaction between PGR2 and PAPPA/IGF may play a role in excess trophoblast invasion in PAS. Furthermore, using previously published single-cell RNA-seq data from early pregnancy samples (21, 26), we confirmed the origin of PAPPA is in interstitial EVTs and syncytiotrophoblasts and PRG2 in interstitial EVTs, supporting this hypothesis (Figure 3b-d). These findings suggest more severe PAS regions are characterised by differential expression of genes which are involved in normal placental trophoblast invasion.

### Reactome pathway analysis of differentially expressed genes

In order to explore the interplay of these genes, we performed reactome pathway analysis of the differentially expressed genes from the spatial transcriptomic data. Pathways involved in degradation of the extracellular matrix (ECM), steroid hormone metabolism and antigen presentation folding assembly and peptide loading of class 1 MHC were upregulated in placenta increta regions (Figure 3e).

Metabolism of steroid hormones pathway was significantly upregulated in PAS regions compared to control. Estrogen and progesterone are the main steroid hormones produced by the placenta (27), with progesterone playing an important role in inhibiting trophoblast migration (28) and estrogen contributing to placental growth through angiogenesis, and upregulating angiogenic factors such as VEGF, Ang-1 and Ang 2 (29). This is in keeping with the finding of increased angiogenesis in PAS as a result of the placenta being implanted over a scarred myometrium (19).

Degradation of the ECM was increased in increta regions compared to accreta. As trophoblasts invade in early pregnancy, degradation of the ECM by metalloproteinases (MMP) is important to allow trophoblast cells to infiltrate the maternal myometrium (30). Following iatrogenic uterine injury, the myometrium demonstrates disorganised and scarred ECM (31), and in PAS, MMPs such as MMP-9 and MMP-2 are increased (32, 33). As PAS is preceded by iatrogenic uterine injury, this suggests increased ECM degradation as a result of myometrial scarring may contribute to deep trophoblast invasion in increta regions.

Antigen presentation folding assembly and peptide loading of class 1 MHC was upregulated in accreta and increta compared to control regions. MHC class 1 peptides are known to play an important role in promoting normal maternal-fetal tolerance by suppressing maternal immune cells (34). Maternal-fetal tolerance is induced by EVT expression of Human leukocyte antigen (HLA) class 1 molecules HLA-G, HLA-E and HLA-C (35). HLA-G is a marker of EVT cells and results in local immune suppression by interacting with decidual natural killer cells and suppressing CD4^+^ T cells (34) and therefore allowing the fetus to escape recognition by the maternal immune system. We found HLA-G was increased in placenta increta regions (Figure 3a), suggesting local immune suppression in these regions. Furthermore, HLA-C is expressed on trophoblast cells in the Beta-2-microglobulin (B2M) form, and is the only HLA on EVT cells which can present paternal antigens and therefore plays an important role in preventing fetal rejection by the maternal immune system (36). We found B2M was increased in PAS regions in our study (see below), similar to previous studies (8, 24). These findings from the spatial transcriptomic data suggest the differentially expressed genes and their associated pathways found to be significant in placenta increta regions play a role in normal placental invasion in the setting of a scarred myometrium and an immunosuppressive environment. In order to investigate this further, we explored the transcription factors involved in regulating these differentially expressed genes in PAS.

### Transcription factors of differentially expressed genes show enhanced invasion capacity in placenta increta regions

The process of normal placental invasion requires coordinated action of various transcription factors, which control the diverse cell groups required for placental invasion including EVT cells and syncytiotrophoblasts (37). In order to explore which transcription factors are active in PAS, we used decoupleR R package (38) to explore transcription factor activity in different PAS regions (Figure 4a).

**Figure 4:**
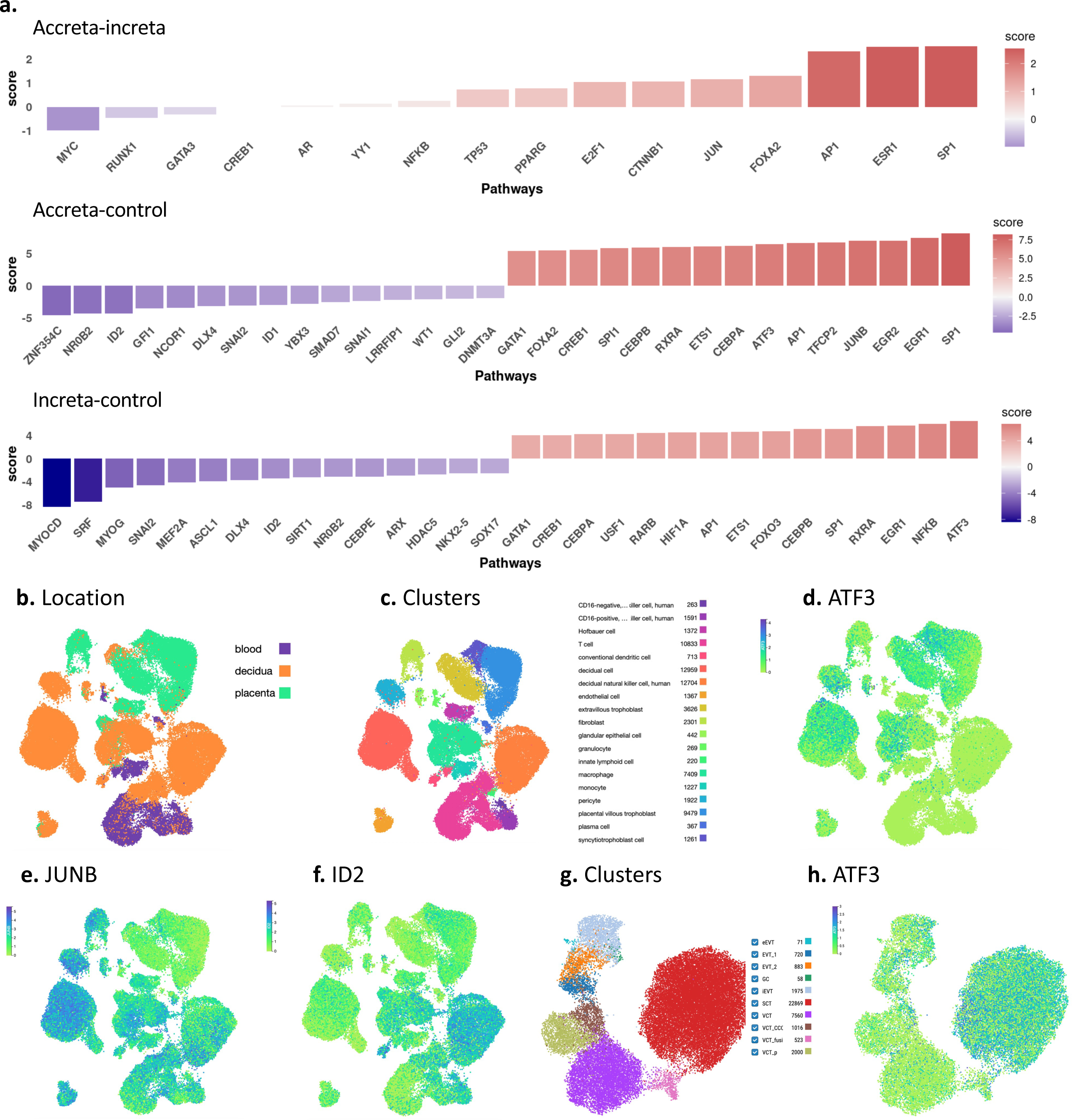
Transcription factors. Figure 4a: Differential transcription factor activity for each comparison, mild-severe, mild-control and severe-control. Transcription factor activities were inferred using the decoupleR R package. **b-f** Single-cell RNA-seq analysis of human first-trimester decidual cells of pregnant women (Vento-Tormo *et a*l 2018, available at https://reproductivecellatlas.org). **b** shows the location and **c** the cell clusters from from 10× Genomics and Smart-seq2 (SS2) scRNA-seq analysis. **d** shows ATF3 expression in extravillous trophoblast (EVT) cells, **e** JUNB expression in multiple cell types in decidua, **d** ID2 predominantly in the decidua in natural killer cells hence confirming our finding of reduced ID2 in PAS which is characterised by reduced NK cells. **g and f;** Spatially resolved single cell data from maternal-fetal interface available as above (hysterectomy specimen); **g** cell clusters from snRNA-seq data, **f** ATF3 also expressed in SCT. *eEVT: endovascular trophoblast cells. GC: placental bed giant cells. iEVT: interstitial extravillous trophoblast. SCT: villous syncytiotrophoblast. VCT: villous cytotrophoblast. VCT_CCC: villous cytotrophoblast cytotrophoblast cell columns*.

Transcription factor activity associated with placenta increta included Specificity Protein 1 (SP1) which accelerates trophoblast cell proliferation and invasion (39) and Estrogen Receptor 1 (ESR1) which, as stated previously, may promote angiogenesis in placenta increta regions. Several transcription factors were also upregulated in all comparisons of PAS regions (accreta and increta) compared to control regions (Early Growth Response (EGR)-1 and 2, ATF3, and AP1).

The most interesting finding from the transcription factor activity analysis was that AP-1 and its subunits, JUNB, JUN and ATF-3 were all factors upregulated in accreta and placenta increta regions compared to control. AP-1 is a transcription factor complex made up of heterodimers of four subfamilies, including Fos, ATF (including ATF3) and JUN (including JUN and JUNB) involved in regulating pro-oncogenic processes such as cellular proliferation and survival, angiogenesis and inflammation in several malignancies including gestational trophoblastic disease, which is characterised by proliferative and invasive trophoblasts (40–43). AP-1 mediates the transcription of programmed death ligand-1 (PD-L1) in lymphoma (44, 45), and PD-L1 was increased in placenta increta regions where AP-1 is highly expressed (see below). In T-cells, AP-1 components induce the transcription of genes that activate or inhibit T-cells (46). This includes activation induced cell death of T-cells, which is triggered by JUN-FOSB heterodimers binding to the promoter of the ligand (CD95L/FAS) of the FAS death receptor, stimulating its expression and inducing cell death during T-cell activation (47).

AP-1 is expressed in EVTs and also regulates invasion and modulates EMT in the placenta (21, 40, 48), and has a number of target genes including matrix metalloproteinase (MMP)-2, MMP-9 and gonadotrophin releasing hormone (GnRH) which are involved in trophoblast invasion (49, 50). AP-1 and NF-κB, another transcription factor upregulated in placenta increta regions (Figure 3a), synchronously regulate several biological processes including immune response (51, 52), and in uterine smooth muscle cells play a synergistic role in parturition through the release of cytokines and regulation of downstream genes such as MMP-9 which causes ECM degradation (53).

ATF3, an AP-1 subunit, which was upregulated in mild and severe PAS regions compared to control, is a common stress response transcription factor (54). ATF-3 was found to be upregulated in severe PAS cases previously (55). Furthermore, ATF3 deficiency has been implicated in defective decidualization and recurrent miscarriage (56). Using previously published single cell RNA-seq data (21, 26), we demonstrate ATF3 is expressed in EVT and syncytiotrophoblast cells (Figure 4b-d). This supports that ATF3 plays a role in trophoblast invasion in PAS.

JUNB plays an important role in establishing normal placentation by regulating gene expression of, for example, MMP-9 and urokinase plasminogen activator (57), which are involved in trophoblast invasion (58). JUNB is recruited by FOSL-1 and forms a transcriptional complex with AP-1, which subsequently controls trophoblast differentiation and invasion (59). We confirmed expression of JUNB in the decidua and in decidual natural killer cells using single cell RNA-seq data (21, 26)(Figure 4e). Therefore it follows that we found JUNB was increased in accreta rather than increta cases, as in increta the decidua is likely more absent/defective than in accreta cases, and increta cases have previously been shown to have reduced natural killer cell populations (15).

In control regions, the expression of the ID (inhibitor differentiation)-2 transcription factor was increased. ID2 is a regulator of trophoblast invasion, and ID2 knockdown results in trophoblast differentiation (60). It is expressed predominantly by decidual natural killer cells, as confirmed by comparison with previous data (Figure 4f). In PAS there is reduced expression of decidual natural killer cells (15), and consequently ID2 expression was reduced in PAS regions.

These results suggest that many of the transcription factors involved in normal trophoblast invasion and promotion of EMT are upregulated in PAS, likely contributing to abnormal placental attachment and deep trophoblast invasion. Several of these transcription factors seem to coordinate with each other to facilitate this process.

### Spatial proteomics suggests the microenvironment in increta regions is characterised by excessive cellular proliferation and activation of mitogen-activated protein kinase pathways in an immunosuppressive local environment

Spatial proteomics was performed on the Nanostring GeoMx DSP (Seattle, WA, USA) and profiled the proteins within each region using GeoMx IO protein panel with nCounter to detect 71 targets. There were 55 differentially expressed proteins between accreta and increta regions (Figure 5a). Overall, differentially expressed proteins were involved in estrogen receptor activation (Figure 5b) in keeping with previous transcriptomic analysis (Figures 3 and 4), inhibition of apoptosis, and immune suppression.

**Figure 5:**
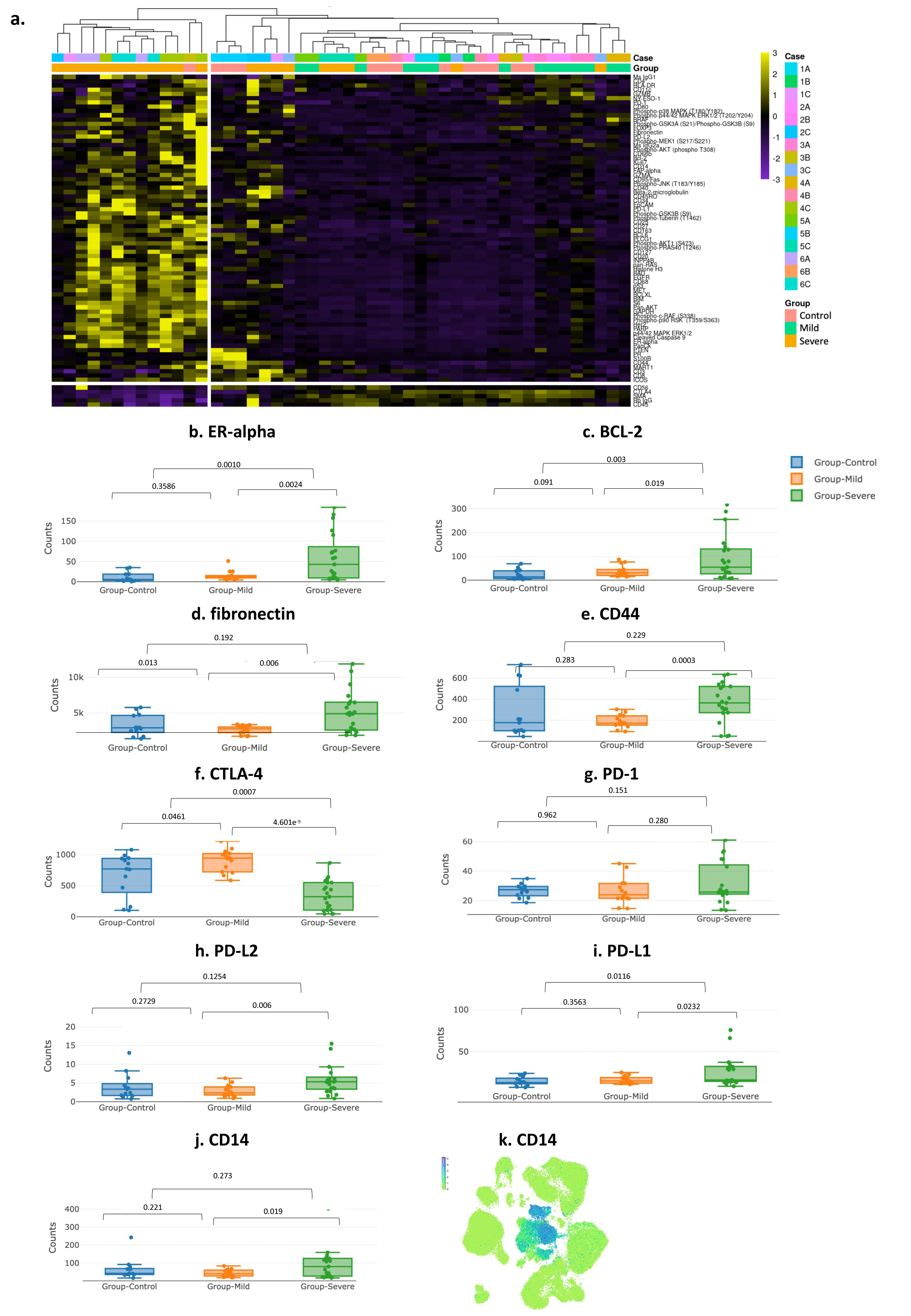
Spatial proteomics. Figure 5a: Heatmap of differentially expressed proteins heatmap showing clustering of controls, accreta and increta cases of the 71 proteins included in the spatial analysis. The heatmap shows the majority of increta (severe) cases clustered together, with proteins involved in immune regulation, anti-apoptosis and estrogen signalling showing increased expression in these regions. **b-j** Differentially expressed proteins - box-plots comparing select proteins between control, accreta (mild) and increta (severe) cases. **k** Single-cell RNA-seq analysis of human first-trimester decidual cells of pregnant women as in figure 4b, CD14 expressed in placental and decidual macrophages

Inhibitors of apoptosis, such as BCL-2, BCLXL and fibronectin (61–63) were upregulated in regions of placenta increta (Figure 5c,d) consistent with previous studies (24, 64). BCL-2 is an important regulator of trophoblast apoptosis in normal pregnancy (65). Fibronectin, an ECM protein, inhibits apoptosis in EVTs by increasing levels of BCL-2 via the PI3K/Akt pathway and therefore increases cell viability and proliferation (63). Fibronectin also plays an essential role in wound healing and repair (66), hence its expression in PAS is not surprising as all PAS cases are preceded by prior uterine injury (18). Interestingly, in a previous PAS study, areas of deep trophoblast invasion were characterised by an abnormally thick layer of fibrinoid at the maternal-fetal interface (19), deposition of which is initiated by fibronectin (20). Fibronectin also increases MMP-9 expression via the AP-1 signalling pathway resulting in ECM degradation (67), which is in keeping with our finding of increased ECM degradation in increta regions in the transcriptomic data (Figure 3e).The combination of increased ER-alpha, BCL-2 and fibronectin expression are likely representative of a more proliferative trophoblastic phenotype in areas of myometrial scarring.

Several mitogen-activated protein kinases (MAPK) pathway proteins, such as p38, Phospho-AKT and extracellular-signal regulated kinase (ERK)-1/2, which are involved in trophoblast differentiation and proliferation (68, 69), were increased in increta regions (Figure 5a). Previously, upregulation of MAPK pathways was found to play a role in trophoblast invasion in PAS through upregulation of MMPs 9 and 2 (49). Furthermore, as mentioned above, fibronectin overexpression results in activation of the PI3K/Akt pathway to inhibit trophoblast apoptosis (63). Therefore, it is possible increased expression of fibronectin in increta regions activates these MAPK pathways, which then stimulates AP-1 transcription factor activity (70). We suggest AP-1 then promotes expression of CD44 and PD-1 and its ligands (see below) resulting in increased trophoblast invasion and immune suppression.

CD44 is expressed in trophoblast cells, plays a role in promoting normal trophoblast invasion (71, 72) and was differentially upregulated in placenta increta regions compared to accreta (Figure 5e). CD44 is associated with increased tumour growth and promotion of EMT (71, 73). AP-1 and NFκB transcription factors, which we found were upregulated in increta regions (Figure 4), have been shown in breast cancer to promote expression of CD44 (74). This suggests these transcription factors may play a role in promoting CD44 and subsequent cell migration and EVT invasion in increta. Interestingly, prominent trophoblast CD44 staining was previously described in a case of PAS, where staining was predominantly seen in the membrane of trophoblasts embedded into the fibrin layer at the fetal-maternal interface (75). CD44 is also a ligand of fibronectin (76), which, as described, was also increased in increta regions.

Spatial proteomics identified several alterations suggestive of an immunosuppressive state in increta regions compared to accreta. Expression of immune checkpoints was altered, with reduced CTLA-4 and increased expression of PD-1 and its ligands PD- L1 and PD-L2 in placenta increta regions compared to accreta regions (Figure 5f-i). Interestingly, we found the CTLA-4 ligand CD80, which is removed from antigen presenting cells by CTLA-4 and is expressed on dendritic cells and monocytes (77, 78), was significantly increased in increta regions. Our findings of reduced CD4^+^ and increased CD25^+^ cells are similar to results from a previous PAS study (14).

PD-1 and its ligand PD-L1 are expressed in syncytiotrophoblast, cytotrophoblasts and EVT (79). Here, they promote an immunosuppressive environment at the maternal-fetal interface (79–81). Among their role in modulating maternal immune cells, PD-1 and PD-L1 expression by trophoblast cells results in macrophage polarization to an M2 phenotype, the dominant macrophage population in the decidua, resulting in increased CD14 macrophages (82). Interestingly, we found CD14 was significantly increased in increta regions (Figure 5j) and confirmed the origin of CD14 as decidual macrophages using single cell RNA-seq data (21, 26) (Figure 5k). Decidual macrophages create an immunosuppressive environment in the decidua by phagocytosing pro-inflammatory cells, expressing anti-inflammatory cytokine such as IL-10 (83, 84) and, as described, PD-1 mediated macrophage polarization results in inhibition of inflammation and immune suppression (82). Alterations in macrophage populations was previously found in PAS using trophoblast cell lines, with increased CD14^+^ reported (85). Several studies have described the role of transcription factor AP-1, which was increased in PAS regions (Figure 4) in regulating PD-1 and its ligand PD-L1 (45, 86). These findings suggest placenta increta regions are characterised by a suppressive immune microenvironment which may allow proliferating trophoblasts to escape the normal regulators of trophoblast invasion. This may result from defective decidualisation as a result of prior uterine injury.

### Enriched ligand-receptor pairs in PAS cases associated with trophoblast invasion in an ITGB1 dependent fashion

Previously, ligand–receptor interactions were found to be enriched in decidual microenvironments during EVT invasion (21). Therefore, we used the BulkSignalR R package (87) to explore ligand-receptor pairs enriched in PAS. Multiple interactions with the ligand/receptor ITGβ1 were active in the PAS regions, particularly in placenta increta (Figure 6a). ITGβ1 increases invasion of EVT cells by activating epithelial– mesenchymal transitions (26, 88), and is expressed in both interstitial EVT and EVT-2 subtypes (21) (Figure 6b). EVT-2s are present at the distal end of the anchoring villous and can differentiate into either interstitial EVTs which invade into the decidual stroma, or into extravascular EVTs which remodel maternal spiral arteries (21). In PAS regions, ITGβ1 interacted with several receptors including fibronectin, A Disintegrin and Metalloprotease domain (ADAM)-9, ADAM12 and the Laminin subunit (LAMC1), each of which are also expressed in various EVT subtypes (Figure 6c-g).

**Figure 6:**
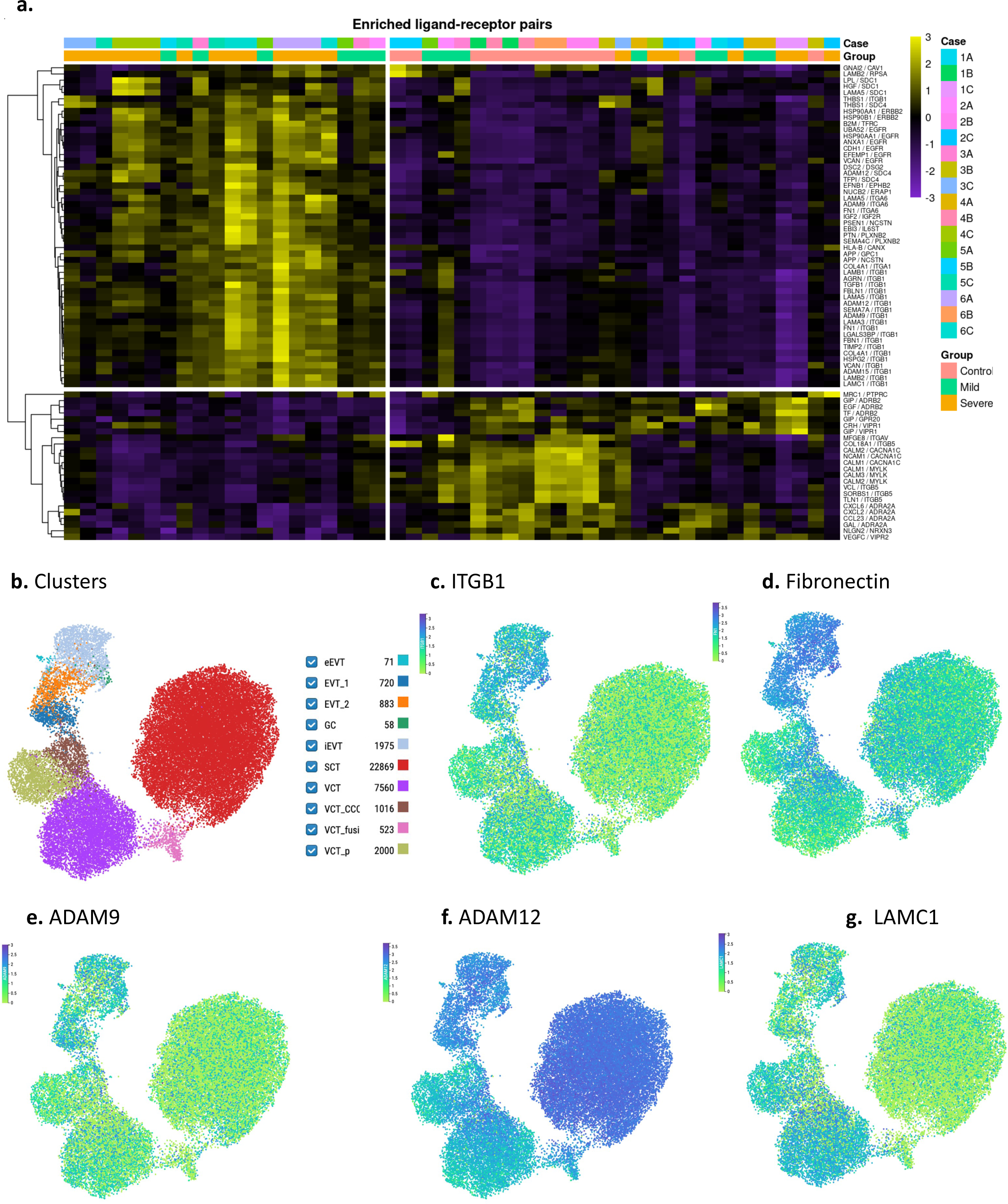
Ligand receptor interaction. Figure 6a Enriched ligand receptor pairs **-** ligand receptor pairs that are active within the dataset using BulkSignalR R package, which were considered significantly active if the q-values were less than or equal to 0.001. **b-g** Spatially resolved single cell data from maternal-fetal interface (hysterectomy specimen, available at https://reproductivecellatlas.org); **b** clusters of cell types from snRNA-seq data, **c** ITGB1 predominantly expressed in intravascular EVT, and EVT 2 subtypes**. d** fibronectin expressed in intravascular EVT and EVT-2, **e-g** other ligands which interact with ITGB1 in PAS regions and their cell expression. *eEVT: endovascular trophoblast cells. GC: placental bed giant cells. iEVT: interstitial extravillous trophoblast. SCT: villous syncytiotrophoblast. VCT: villous cytotrophoblast. VCT_CCC: villous cytotrophoblast cytotrophoblast cell columns*.

ITGβ1 binding to fibronectin is in keeping with the proteomic data where fibronectin was differentially expressed in increta regions (Figure 5). We confirmed the origin of ITGβ1 and fibronectin as iEVT and other EVT subtypes using single cell-RNA seq data (21) (Figure 6c,d). ITGβ1 and fibronectin signalling has been shown to promote cell migration (89), as does fibronectin and ITGA6 binding (90), which was also upregulated in increta regions. Furthermore, fibronectin and ITGβ1 ligand interaction inhibits T cell activation and upregulates macrophages (91), consistent with our findings in increta cases in the immunohistochemistry (figure 2) and protein data (figure 5).

ADAM-9, which is predominately expressed in interstitial EVTs (21) (Figure 6e), is involved in mediating cell adhesion and migration and is pro-angiogenic in tumours through ECM remodelling (92, 93). ADAM12 is highly expressed in the placenta (Figure 6f) and promotes placental development by increasing trophoblast migration and invasion by degradation of ECM (94). ADAM12 is specifically expressed in invasive ECM degrading EVTs (94), in keeping with our finding of increased ECM degradation in increta regions (Figure 3e).

Laminins are a family of ECM proteins that are involved in several processes including cell adhesion, differentiation and migration (49), which are important for establishing normal placentation in early pregnancy (95). Moreover, a previous PAS study found trophoblast cells over expressed laminins (10) and a mouse model found lack of laminin resulted in defective and shallow trophoblast invasion (96). This further suggests placenta increta regions are characterised by upregulation of the normal processes of placental invasion, which may be driven by enriched ligand receptor interactions with ITGβ1 which is a ligand of fibronectin.

## Discussion

Using an analytical approach consisting of exploration of transcriptomic alterations and their associated pathways and transcription factors, followed by spatial proteomics and ligand receptor interactions, we found regions of placenta increta are characterised by overexpression of genes involved in normal trophoblast invasion and EMT. Upregulated pathways in increta were related to ECM degradation and immune suppression, with AP-1 signalling and its subunits emerging as important transcription factors in PAS. The protein data suggests AP-1 may be activated by phosphorylation of MAPK pathways, with AP-1 signalling then activating CD44 and PD-L1 expression, resulting in increased trophoblast invasion and local immunosuppression. Fibronectin expression appears to be a central factor in abnormal placental attachment in PAS through several processes such as inhibition of apoptosis and interacting with proteins such as CD44 to promote cell proliferation. Furthermore, ligand receptor interactions with fibronectin and ITGβ1 may cause degradation of ECM and immune suppression, further contributing to abnormal trophoblast invasion in PAS. The findings from this study and possible inferred pathways from these results in PAS are summarised in Figure 7.

**Figure 7:**
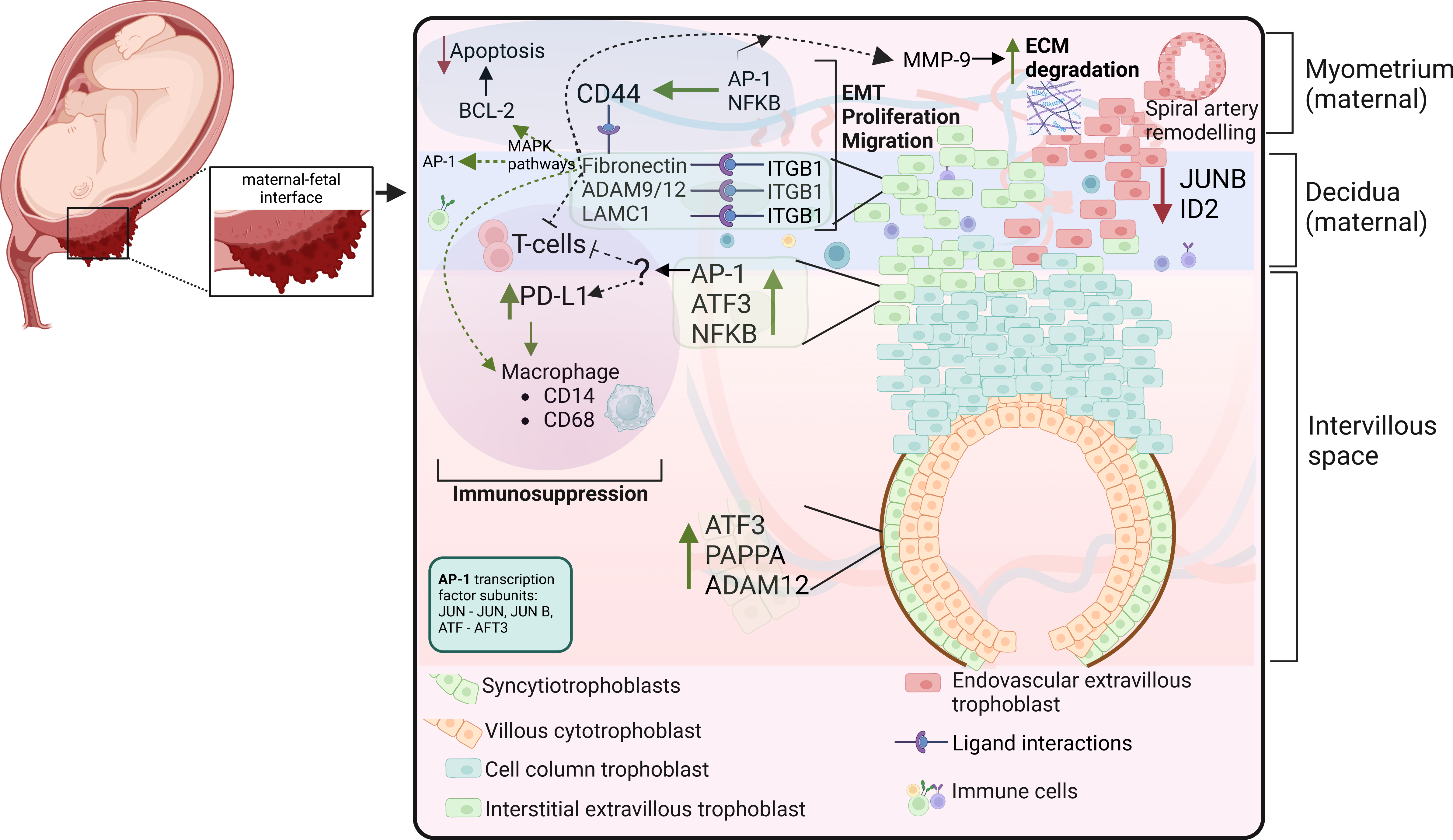
Summary of results and inferred biological explanations contributing to PAS. Figure 7 summarises the results from the analytical approach utilised in this study, consisting of spatial transcriptomics, pathway analysis, transcription factors, spatial proteomics and enriched ligand receptor interactions. The dotted lines and arrows represent inferred biological explanations which are possible downstream events of the findings shown in this study that may contribute to abnormal placental attachment in PAS. Green lines and arrows represent upregulation or increased expression of genes and proteins, while red represents downregulated or reduced expression.

Our study found increta regions are characterised by expression of genes and proteins associated with an overly invasive EVT phenotype. For example, in increta regions there was increased expression of genes, such as NF-κB2, involved in EVT invasion (Figure 3a), as well as upregulation of transcription factors which promote EVT invasion and EMT such as AP-1 and its subunits, and reduced transcription factors which negatively regulate EVT invasion such as ID2 (Figure 4). Enriched ligand interactions in increta regions were predominantly with ITGβ1, expressed in EVTs (Figure 6b), and other promotors of cell invasion and adhesion and ECM degradation such as fibronectin, ADAMs and laminins (Figure 6c-f). In turn, fibronectin is a ligand of CD44 which also promotes cell invasion and adhesion in trophoblasts and is upregulated by the transcription factors AP-1 and NF-κB (63, 71, 72, 74). Fibronectin, as an ECM protein, initiates fibrinoid deposition along with other ECM proteins in normal placentas to anchor the placenta to the myometrium (20), and as described, a previous PAS study reported excessive deposition of fibrinoid at the maternal-fetal interface (19). Fibronectin subsequently inhibits trophoblast apoptosis and cell proliferation via the PI3K/Akt pathway (63). Therefore, fibronectin expression and subsequent excessive fibrinoid deposition and cell proliferation is likely an important contributor to the abnormal placental adherence seen in PAS.

Furthermore, our results suggest there is local immune suppression in increta regions. This may be driven by increased AP-1 signalling, resulting in increased expression of PD-L1 and PD-L2 (Figure 5b) (45), which promotes macrophage polarization to an M2 type in the decidua and subsequent immunosuppression (82–84). Fibronectin and ITGβ1 also inhibit T cell activation and upregulate macrophages (91). This suggests AP-1 signalling of PD-1 and its ligands creates an immune suppressed state in PAS, which may allow EVTs to escape some of the normal immune regulators of EVT invasion.

These biological changes are likely secondary to the placenta implanting over a previously scarred myometrium, as almost all cases of PAS are preceded by an iatrogenic uterine injury, usually a Caesarean section (5, 18). Following iatrogenic injury, the myometrium fails to fully heal, demonstrating impaired wound healing in the form of elastosis, myofiber disarray and scar formation (31, 97). Therefore, it is likely many of the biological changes associated with an invasive EVT phenotype occur secondary to this myometrial scarring and the resultant defective decidua (24). This is supported by our finding of increased ECM degradation in increta regions (Figure 3e). A possible mechanism for this in PAS based on our findings may be EVTs gaining deeper access to and remodelling maternal spiral arteries as a result of local immune suppression and increased cell proliferation secondary to prior iatrogenic uterine injury. Subsequently there is increased expression of fibronectin as described above from chronic high velocity flow from these remodelled spiral arteries, which upregulates MMP-9 expression via the AP-1 signalling pathway and increases ECM degradation (67). Furthermore, ITGβ1 ligand interactions with ADAM-12 and fibronectin contributes to ECM remodelling. Subsequently, ECM degradation facilitates increased trophoblast invasion by removing the physical barriers in the myometrium to EVT invasion. Taken together with an increased invasive EVT phenotype in an immunosuppressive environment, this results in the abnormal placental attachment and deep trophoblast invasion seen in PAS.

This study has a number of strengths and limitations. This study represents a well classified cohort of PAS cases obtained prospectively who were all cared for by an established PAS MDT. Using a spatial approach, we were able to select specific regions at the maternal-fetal interface to study, which is particularly important in PAS as this is the area where abnormal placental adherence or invasion occurs. We suggest future studies investigating PAS utilise spatial approaches as the maternal-fetal interface and alterations here are key to understanding the underlying mechanisms leading to abnormal placenta attachment. This study is limited by the small number of cases included. Furthermore, the number of transcripts and proteins analysed was relatively small as a result of the current limitations of spatial proteomic technology. The findings presented here, in particular the inferred biological downstream events of the findings shown in this study, will require validation in future studies.

In conclusion, we found several processes may contribute to abnormal placental adherence in PAS. Fibronectin appears to play a central role in this process, through interaction with CD44, ITGβ1, and inhibiting apoptosis, which promotes local immune suppression and increases EVT invasion. Disorganised and increased degradation of ECM, likely as a result of myometrial injury, further contributes to EVTs gaining deeper access to the myometrium. These changes are likely secondary to defective decidualisation as a result of previous iatrogenic myometrial injury. Therefore, this study provides further insights into the underlying processes leading to PAS using a spatial multi-omics approach.

## Materials and Methods

### Study population

Data was prospectively obtained between January 2018 – October 2022. Ethical approval was obtained from the National Maternity Hospital, Dublin (EC30.2018) and participants provided written, informed consent. Inclusion criteria were as follows: women who had suspected PAS based on ultrasound assessment by a fetal medicine specialist with features as previously described (98), and who at the time of laparotomy had intraoperative clinical features of PAS as per the FIGO classification (5), and with final histopathological confirmation as examined by a perinatal histopathologist (> 10 years of experience) as previously described (supplementary figure 1)(7). Cases were classified as either placenta accreta or placenta increta as follows: placenta accreta was diagnosed when placental villi were seen in direct contact with the superficial myometrium and no intervening decidua, and as placenta increta where placental villi were seen deep within the muscular fibres and/or in the lumen of the deep uterine vasculature (Supplementary figure 1). Controls were defined as patients with a prior Caesarean section and placenta previa diagnosed on ultrasound, with spontaneous placental separation at the time of Caesarean section. Women who met the following criteria were offered to participate as controls: sonographic finding of placenta previa, defined as the placenta completely covering the internal cervical os on transvaginal ultrasound beyond 20 weeks’ gestation (99), and history of at least one prior Caesarean section, and no sonographic evidence of PAS and spontaneous placental separation at the time of Caesarean section. All participants were cared for by a PAS multi-disciplinary team (100).

### Immunohistochemistry

For the immunohistochemistry, 41 participants met inclusion criteria (n=35 cases, n=6 controls). Cases and controls had a similar median age, parity and number of prior Caesarean sections (supplementary materials table). Four antibodies were chosen to establish their expression pattern in placental samples: CD3^+^, cytokeratin (CK)7, CD8^+^, and CD4^+^ (supplementary materials table 2). For each selected case, 4µm formalin-fixed, paraffin-embedded, placental sections were immunohistochemically stained for each of the aforementioned biomarker. Briefly, the slides were stained separately for CK7, CD3^+^, CD4^+^, and CD8^+^ on the Ventana Benchmark Ultra system with the Optiview DAB IHC Detection Kit. The stained slides were then scanned using an Aperio AT2 digital slide scanner (Leica Biosystems Ltd.) at 20X magnification. The uterine-placental interface of the scanned images were annotated using ImageScope software (Leica Biosystems Ltd.). The Aperio Nuclear Algorithm v9 and Aperio Colour Deconvolution Algorithm v9 (Leica Biosystems Ltd.), were trained to detect positive staining in the annotated images. The expression of CD3^+^, CD4^+^, and CD8^+^ was calculated as the number of positively stained cells per mm^2^. For CK7 expression was determined as area of positive staining per mm^2^.

### Tissue processing: spatial transcriptomics and proteomics

Six cases that were included in the immunohistochemistry were selected for spatial transcriptomic and proteomic analysis. Patient demographics are shown in Table 1, and all participants underwent Caesarean hysterectomy. For each case, three separate areas for analysis were selected; an area of placenta accreta (a), an area of placenta increta (c) and an internal control (b), consisting of myometrial muscle away from the fetal-maternal interface. For case 6, there was no area of placenta accreta identified, hence two regions with placenta increta were selected and paraffin embedded. Areas for analysis (placenta accreta, placenta increta and control) were identified by examining H&E slides prepared with FFPE tissue. These areas were then biopsied from the FFPE tissue blocks, and formatted into a single block for each case. A 4um section from each formatted block was prepared with H&E to ensure suitability for the modified tissue microarray (TMA). Each formatted block was sectioned onto the same slide to create a TMA with 18 samples (3 from each case) (Figure 1, supplementary materials figure 2) which were baked at 100 degrees Celcius for 60 minutes. The slide was processed for spatial transcriptomics and proteomics analysis according to the GeoMx DSP protocol (101), provided by Nanostring (Seattle, WA, USA). Using the GeoMx Digital Spatial Profiling, the modified TMA slide was examined and regions of interest (ROI) were selected. A total of 48 ROIs were selected for analysis, which consisted of the maternal myometrium at the maternal-fetal interface (supplementary materials figure 2). Using pre-built panels, the RNAs within each region were profiled using GeoMx Human Whole Transcriptome Atlas to detect approximately 12000 targets, while protein within each region was profiled using GeoMx IO protein panel with nCounter to detect 71 targets.

### Spatial RNA expression analysis

Data processing was performed following the Nanostring’s guidelines for GeoMx-NGS RNA expression analysis using the recommended tools (102), viz-NanoStringNCTools, GeoMxWorkflows and GeoMxTools packages. Briefly, for initial data pre-processing, quality control was performed for each segment ROI/AO to assess the quality of sequencing and tissue sampling. Here, the quality threshold was set at minSegmentReads = 1000, percentTrimmed = 80, percentStitched = 80, percentAligned = 80, percentSaturation = 50, minNegativeCount = 5, maxNTCCount = 6000, minNuclei = 100, and minArea = 5000. Interestingly, 47 ROIs passed the quality assessment but the only 1 ROI not passing the quality control assessment was excluded from further analysis (supplementary materials figure 2). Next, using a principal component analysis (PCA), case replicate clustering was performed to assess and visualize the homogeneity within and/or heterogeneity between cases based on gene expression; replication ROIs within cases were found to cluster well together and apart from those of other cases (supplementary materials figure 3). Furthermore, a probe-level quality control was performed using the default parameters. Finally, for gene-level quality control, a limit of quantification (LOQ) threshold was defined using the default standard deviation and minLOQ of 2. Setting the minimum gene detection rate (the proportion of ROIs where each gene is detected) at 10%, a total of 11,795 genes passed the quality control filtering and were brought forward for further analysis. After quality control processing, data normalization was performed using the background normalization method from the GeoMx ‘Normalize’ function, with the default parameters but method set to ‘neg_norm’. Following data pre-processing, differential expression analysis was performed by fitting a linear mixed-effect model without a random slope. The differential expression analysis was performed in 3 pairs-control vs placenta accreta regions, control vs placenta increta regions, and accreta vs increta regions. Significantly differentially expressed genes (DEG) were those for which p-values <0.05 and minimum log2 fold change (FC) of 0.5 (supplementary Tables DEGS).

### Functional annotation

Gene set enrichment analysis (GSEA) of the DEGS were performed using the ClusterProfiler (103) and fgsea R packages (104). Significantly enriched pathways were those for which p-values < 0.05.

### Transcription factor network analysis

Transcription factor activities were inferred using the decoupleR R package (38) which is based on a univariate linear model (ULM). The model allows the determination of active or inactive transcription factors based on the gene expression pattern in the dataset of interest. Essentially, this is achieved by querying the collection of transcriptional regulatory interactions (CollecTRI) which consists of comprehensively curated TFs and their corresponding transcriptional targets obtained from 12 curated resources and available in the Omnipath database. So, for each sample in the dataset and each transcriptional factor in the CollecTRI network (TF-genes), decoupleR fits a ULM and scores each TF while also giving account of the regulatory directionality (active or inactive). In our own case, we seek to understand differential changes in TF activities between cases, therefore, decoupleR was run against the log2FC values of the significant DEGs from the 3 comparison pairs-control vs accreta regions, control vs increta regions, and accreta vs increta regions. The TFs were ranked based on the enrichment scores and the top 30 differentially active and inactive TFs were taken for each of the compared groups.

### Ligand-receptor interaction analysis

The BulkSignalR R package (87) was utilized to depict possible ligand-receptor interactions. This method works by taking into consideration not only the expression of ligands and receptors but also the expression of the target genes of the downstream signalling pathway. Briefly, each dataset is decomposed into a triple of ligand-receptor-pathway (LRPw) such that for each ligand-receptor pair (LRP) gene expressed, the expression of the downstream pathway target genes must be correlated and statistically significant before considering that LRP as being active. Essentially, this tool scans the ligand-receptor database for information about LRPs, and queries the KEGG, Gene Ontology Biological Process and Reactome pathways to delineate downstream pathways. Target genes are inferred from the reference network topology as those genes reachable from a receptor in a pathway by the directional edge annotated as “controls the expression of”. LRPs were considered significantly active if the q-values were less than or equal to 0.001.

### Analysis of publicly available single-cell RNA sequencing and transcriptomic spatial data from early pregnancy

In order to determine the origin of several transcription factors (ATF-3, ID2, JUNB, PAPPA, PGR2) and ligands (ITGβ1) of interest in PAS regions, we analysed two public available datasets; one which reported single-cell RNA sequencing data of human first trimester (6-14 gestational weeks) decidual cells (26) and another which reported spatially resolved single cell multiomic characterisation of the maternal fetal interface 6-9 gestational weeks (P13 used as the fetal-maternal interface represented as specimen of hysterectomy, both datasets available here https://reproductivecellatlas.org.(21)).

## Acknowledgements

This research was funded by Placenta Accreta Ireland, a patient advocacy group, the Medical Fund at the National Maternity Hospital and the National Maternity Hospital Foundation, Dublin, Ireland, and Science Foundation Ireland (SFI) under Grant Number 18/SPP/3522.

The funders played no role in the study design, analysis or results.

Figure 1 and 7 created using Biorender (biorender.com)

---

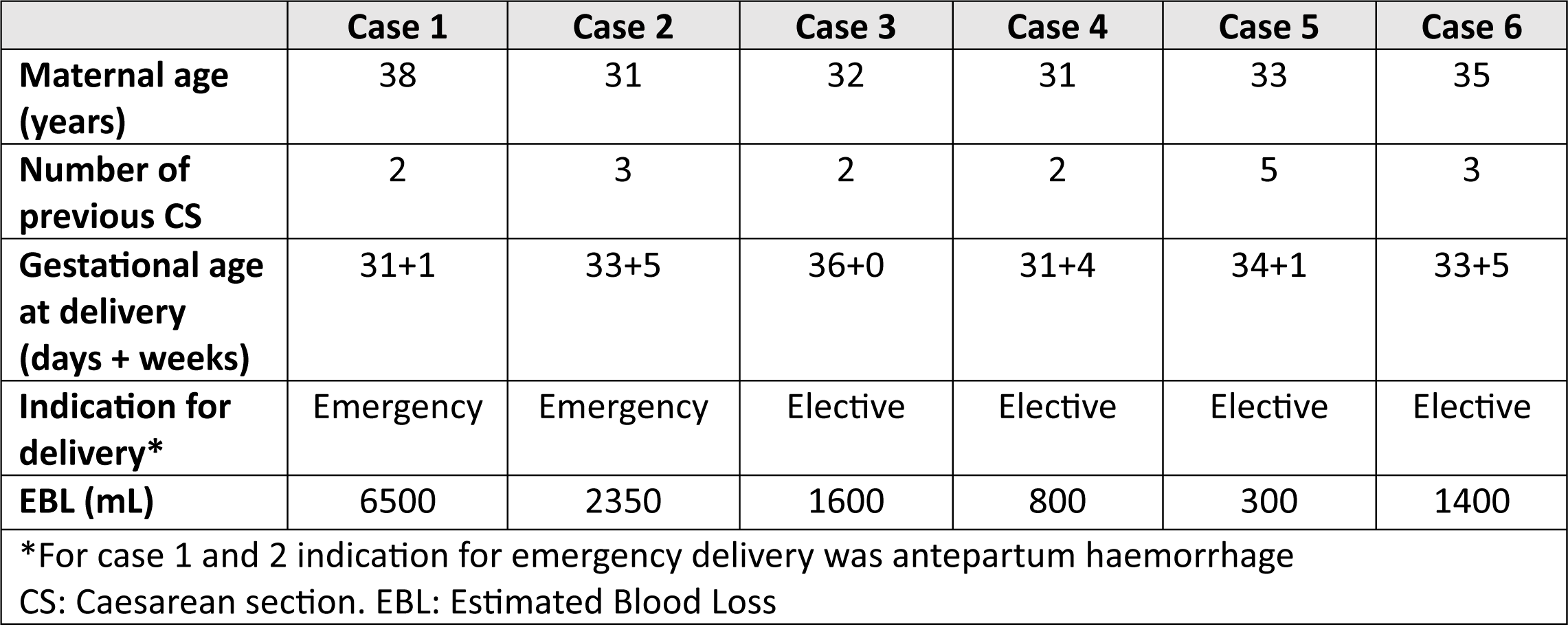

