## Supplementary figures and tables for "Spatial proteomics and transcriptomics of placenta accreta spectrum"

### Supplementary materials
### Figure 1: Histopathological case confirmation and slide selection

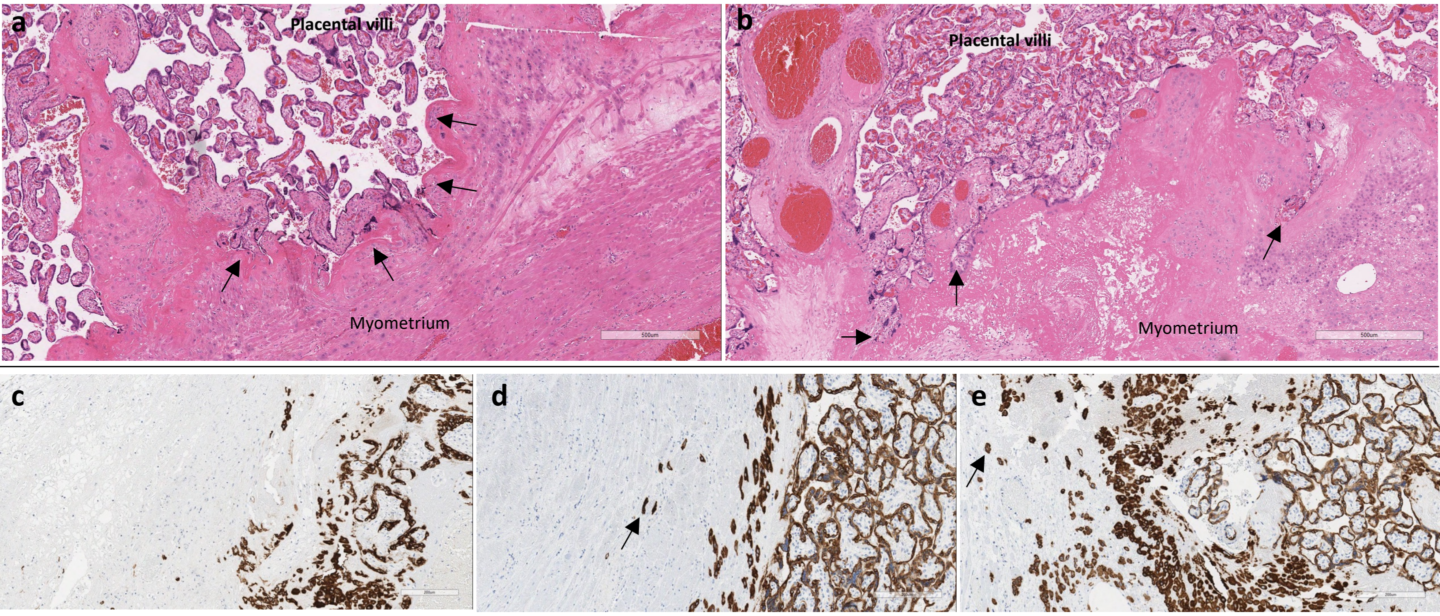


Figure 1a and 1b shows two examples of selected H&E stained slides at 500um magnification. Image **a** shows a case of placenta accreta, where placental villi are in direct contact with myometrium with no intervening decidua. Black arrows show area of normal contour of maternal-fetal interface with a thin layer of fibrin (darker pink). Image **b** shows a case of placenta increta, trophoblast cells are seen invading deep into the myometrium (arrows). The maternal-fetal interface is less clearly defined. Image **c, d** and **e** show cytokeratin-7 (CK-7) stained slides at 200um magnification. CK-7 stains for fetal trophoblast cells. Image **c** shows an example of placenta previa, placenta accreta and placenta increta. In **c,** a normal level of trophoblast invasion is seen, while in **d** trophoblast invasion beyond a deficient basal plate and decidua into the myometrium is present (arrow), with deep invasion seen in **e** (arrows).

#### Supplementary materials: Figure 2
#### Workflow of tissue preparation and region of interest selection

##### 2a: Modified tissue microarray

Six PAS cases with three tissue areas were selected (N=18). Tissue areas were selected as follows: area of placenta accreta (Area B), area of placenta increta (Area A) and an internal control, selected as an area of myometrium away from the fetal-maternal interface (Area C). H&E stained slides were examined to identify the three areas of interest. A 3mm bunch biopsy was used to obtain the samples from tissue blocks and create a modified tissue microarray (TMA), with cases labelled as in image 1a.

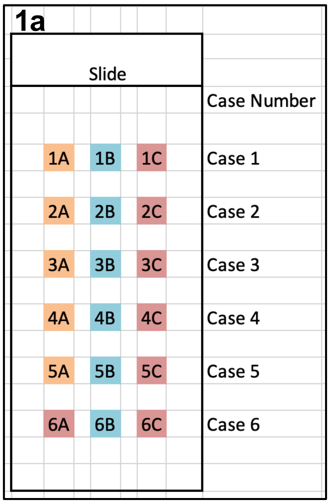


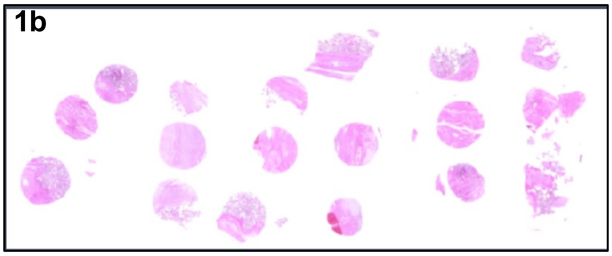


**1b:** H&E stained slide of TMA slide to confirm each area was correctly sampled.

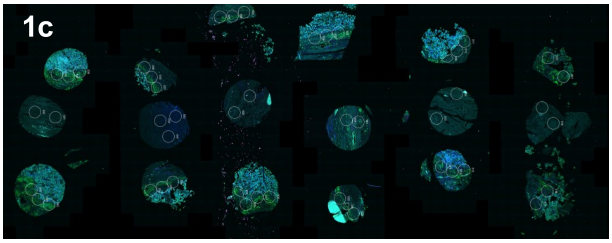


**1c:** TMA scanned using GeoMx DSP

##### 2b: Geometric Segmentation

Regions of interest (ROIs) were selected using the GeoMx DSP control centre (Version 2.5.1.145). The corresponding H&E TMA was reviewed when selecting the ROIs (1,2,3). Images d,e,f show examples of a control (e), accreta (d) and increta (f) case. Staining with antibodies against panCK (blue), DNA (green), CD45 (yellow) and CD56 (Red). Images at 300um magnification.

**a. placenta accreta** **b. Control** **c. placenta increta**

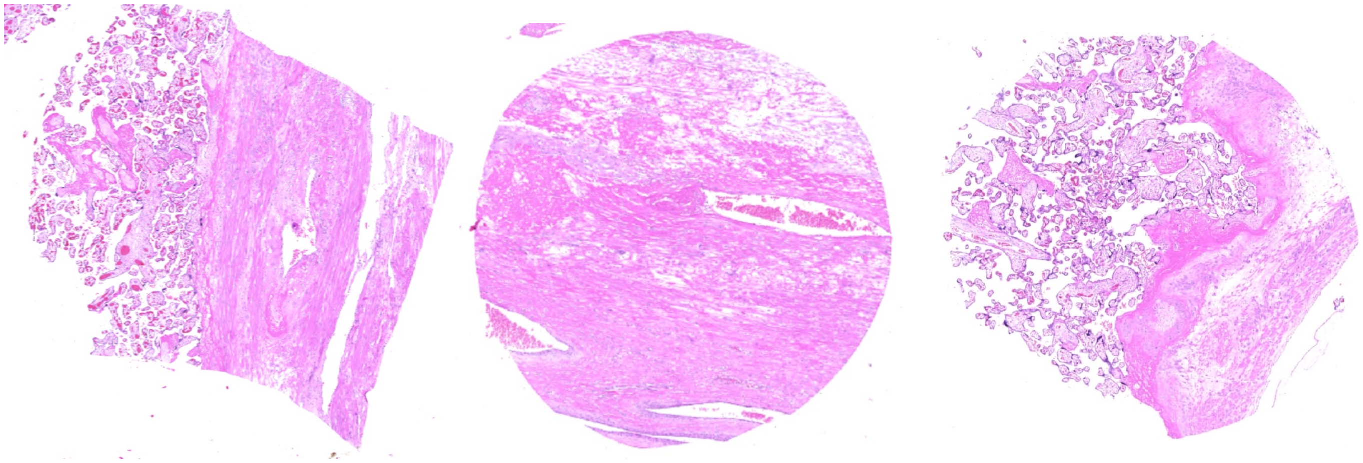

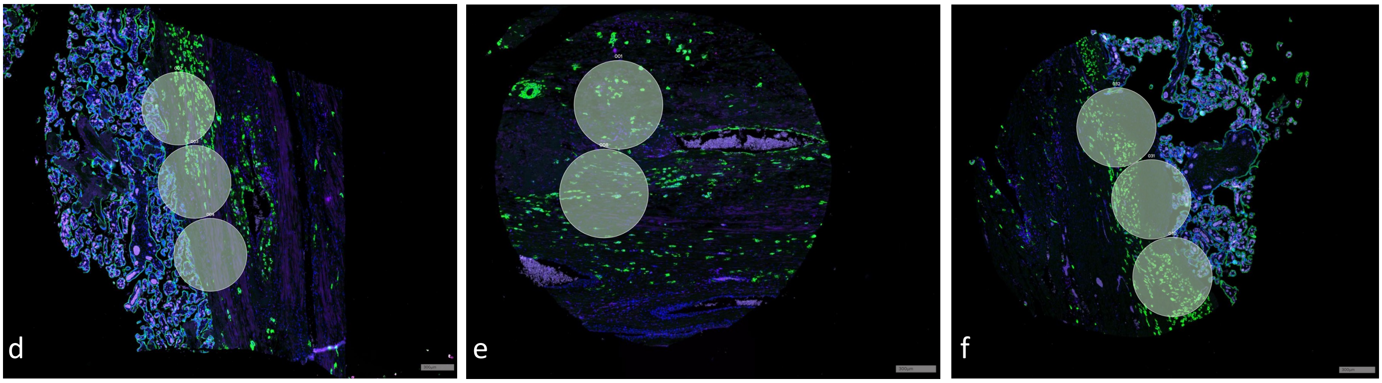

**Supplementary materials:**
**Figure 3 Quality control**

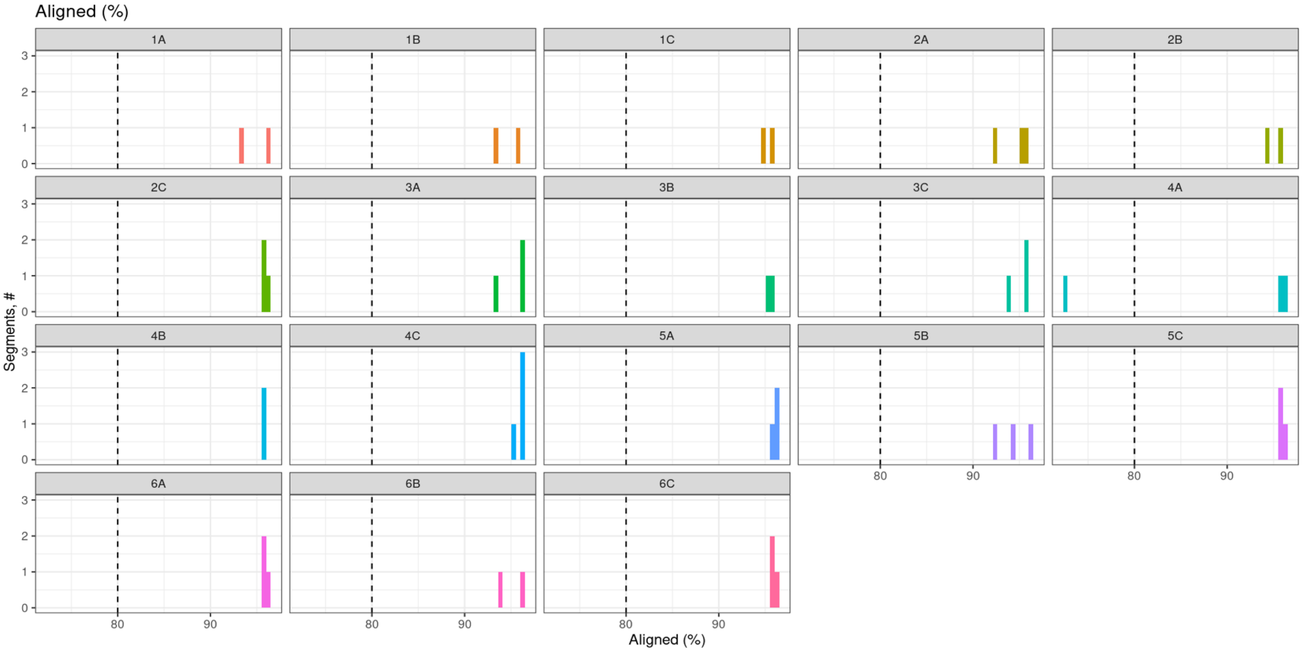


**Figure 3** shows the quality control target assessment for each case, with only one region in 4a not passing the quality control assessment.

**Supplementary materials:**
**Figure 4 Case/Technical replicate clustering**

a.

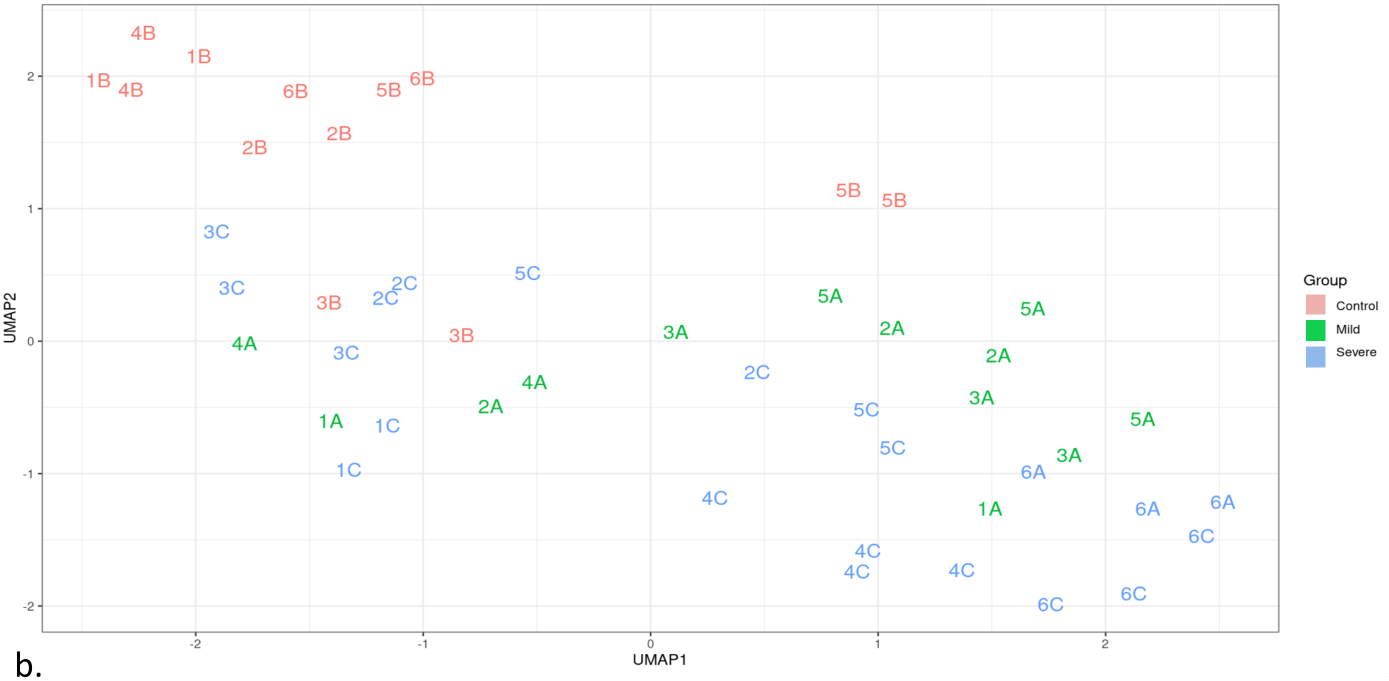


b.

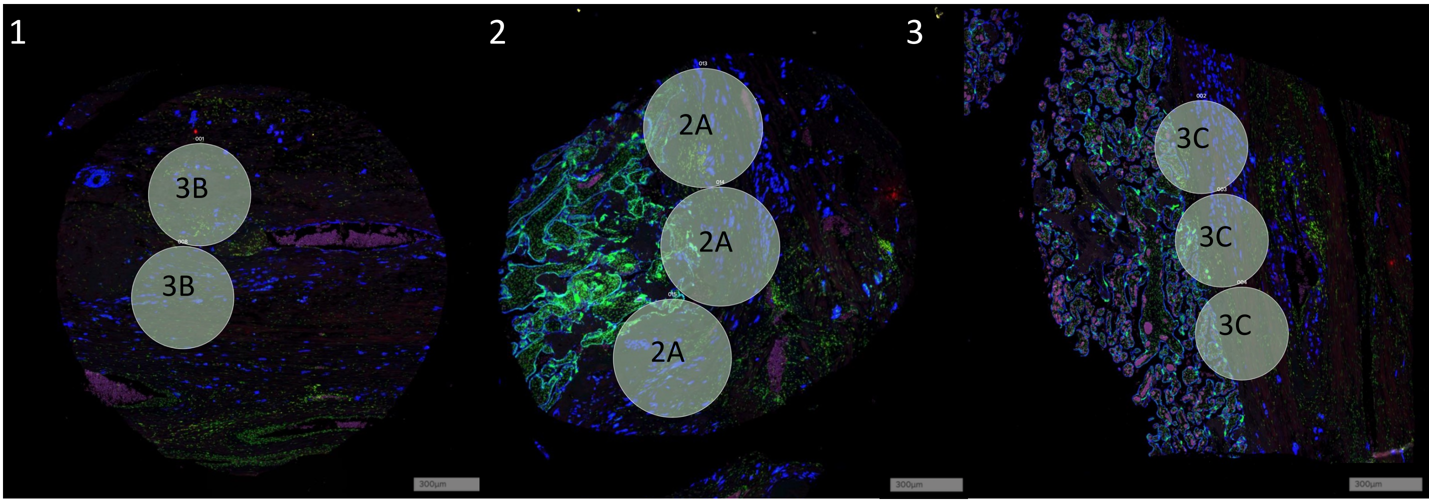


**Figure 4**: ROIs within each tissue sample were compared to assess for heterogeneity within each sample. **4a** shows good clustering of replicate ROIs within each sample, meaning further analysis pooled the ROIs from each case together. **4b** shows an example of a control (1), placenta accreta (mild) (2) and placenta increta (severe) (3) case. Staining with antibodies against panCK (blue), DNA (green), CD45 (yellow) and CD56 (Red).

*ROI: Region of Interest*

| Table S1: Demographics and Pregnancy outcome for immunohistochemistry | | | |
|---|---|---|---|
| **Participant demographics** | **Total N=41** | **Cases N=35** | **Controls N=6** |
| Age | 38 (34.5-40.0) | 38 (34-40) | 37.5 (33.7-39.7) |
| BMI (kg/m2) | 25.4 (22.7 – 29.7) | 26.5 (23.2 – 31.7) | 23.3 (21.9 – 25.5) |
| Parity | 2 (1-2.5) | 2 (1-3) | 2 (1-2.5) |
| Number of previous CS | 1 (1-2) | 1 (1-2) | 1 (1-2.5) |
| Risk factors *(history of)* N (%) | | | |
| *ERPC* | 14 (34.1) | 12 (34.3) | 2 (33.3) |
| *MROP* | 10 (24.3) | 8 (22.8) | 2 (33.3) |
| *Placenta previa* | 12 (29.3) | 10 (28.6) | 2 (33.3) |
| *ART* | 5 (12.2) | 5 (12.2) | 0 |
| **Pregnancy outcome** | | | |
| Type of delivery N (%) | | | |
| Emergency | 11 (26.8) | 9 (25.7) | 2 (33.3) |
| Elective | 30 (73.2) | 26 (74.3) | 4 (66.7) |
| Gestational age at delivery | 35 (28-38) | 34+5 (28-36) | 35 (33-37) |
| Surgical procedure N (%) | | | |
| *Hysterectomy* | 25 (61) | 25 (61) | 0 |
| Estimated Blood Loss (ml) | 1400 (300-8250) | 1500 (800-3500) | 600 (400-1141) |
| Blood transfusion | | | |
| *Received RCC N (%)* | 15 (36.6) | 14 (40) | 1 (16.7) |
| *Units of RCC transfused* | 4 (2-6) | 4 (2-6) | 4 |
| Histological grade N (%) | | | |
| Placenta accreta | 18 (47.3) | 18 (47.3) | |
| Placenta increta | 17 (48.5) | 17 (48.5) | |

Values presented as median (IQR) unless otherwise stated.

*BMI: Body Mass Index. CS: Caesarean Section. ERPC: Evacuation of Retained Products of Conception. MROP: Manual Removal of Placenta. ART: Artificial Reproductive Technique. RCC: Red Cell Concentrate.*

| Table S2: Antibodies for immunohistochemistry analysis | | | |
|---|---|---|---|
| | Brand | Clone | Category Number |
| CD3 | Roche Diagnostics (Ventana) | 2GV6 Rabbit Monoclonal | 05278422001 |
| CK7 | Roche Diagnostics (Ventana) | SP52 Rabbit Monoclonal Primary Antibody | 05986818001 |
| CD8 | Dako Agilent | C8/144B, Mouse Dilution: 1:25 | M7103 |
| CD4 | Roche Diagnostics (Ventana) | SP35, Rabbit Dilution | 05552737001 |
